# Nucleotide metabolism constrains prime editing in hematopoietic stem and progenitor cells

**DOI:** 10.1101/2023.10.22.563434

**Authors:** Sébastien Levesque, Archana Verma, Daniel E. Bauer

## Abstract

Therapeutic prime editing of hematopoietic stem and progenitor cells (HSPCs) holds great potential to remedy blood disorders. Since quiescent cells have low nucleotide levels and resist retroviral infection, we hypothesized that nucleotide metabolism could limit reverse transcription mediated prime editing in HSPCs. We demonstrate that deoxynucleoside supplementation and Vpx-mediated degradation of SAMHD1 improve prime editing efficiency in HSPCs, especially when coupled with editing approaches that evade mismatch repair.

## MAIN

Prime editing (PE) relies on a Cas9 nickase fused to a reverse transcriptase along with an extended prime editing guide RNA (pegRNA) to introduce to living cells templated genomic modifications, including point mutations, short insertions and deletions (indels), and longer sequence replacements^1–3^. While recent advances have improved the technology, efficient prime editing remains challenging in primary hematopoietic cells. One key difference between cancer cell lines and primary hematopoietic cells is the concentration of nucleotides available for reverse transcription. Nondividing cells typically have orders of magnitude lower nucleotide levels as compared to dividing cells^4–6^. SAMHD1 is a triphosphohydrolase enzyme that depletes deoxyribonucleoside triphosphates (dNTPs) and acts as an antiviral factor to restrict HIV-1 infection^7–9^. Since SAMHD1 expression markedly increases in hematopoietic stem and progenitor cells (HSPCs) after cytokine culture^10^, we hypothesized that modulating the nucleotide metabolism could enhance reverse transcription and thus prime editing (**Fig. 1a**).

**Figure 1.**
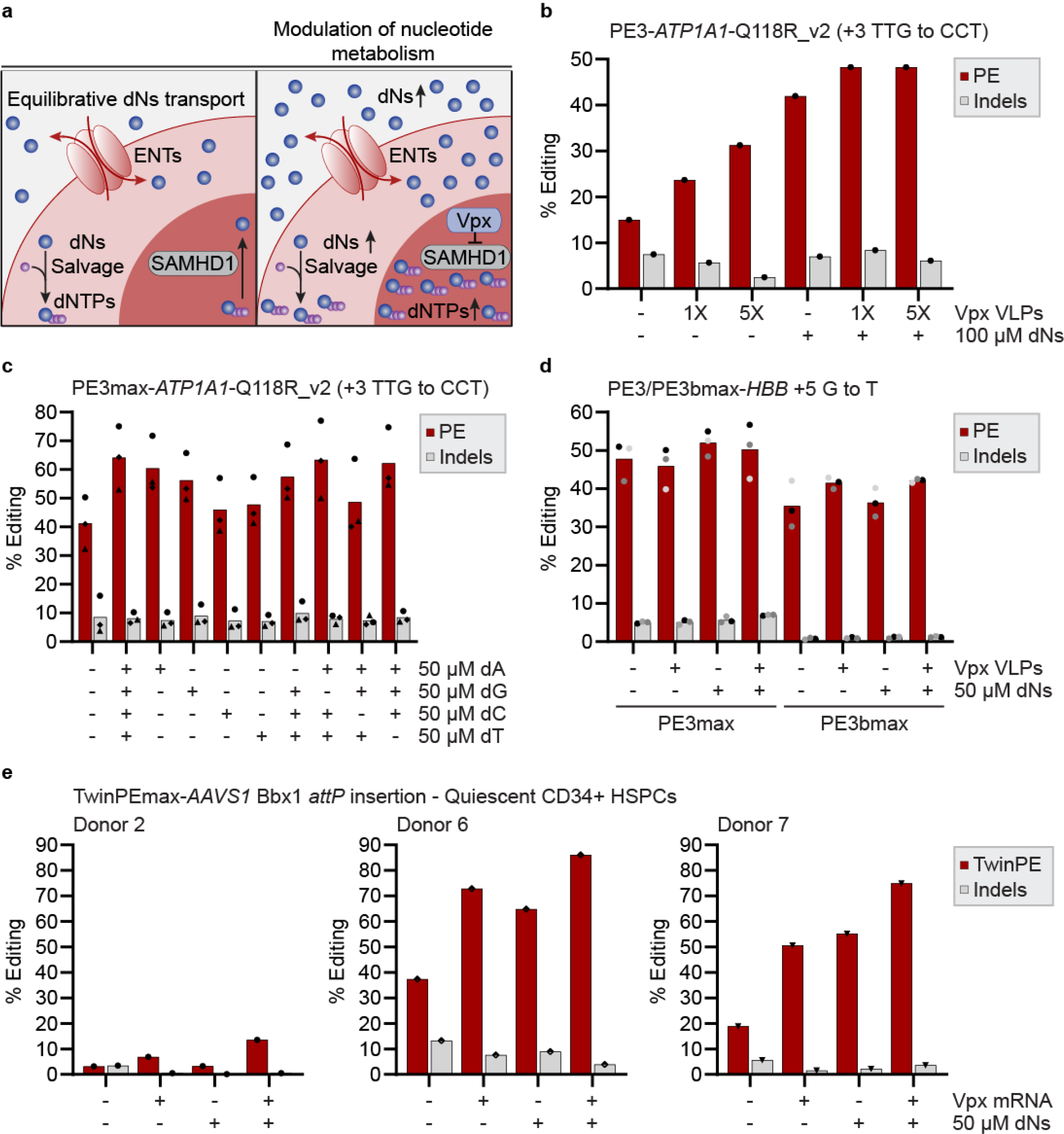
Vpx and deoxynucleoside (dN) supplementation enhance prime editing in hematopoietic stem and progenitor cells. (**a**) Schematic representation of the strategies used for the modulation of nucleotide metabolism to increase the concentration of dNTPs available for reverse transcription and prime editing. (**b**) PE and indels quantification as determined by BEAT and TIDE analysis from Sanger sequencing. HSPCs were thawed and cultured in the presence or absence of 100 µM dNs for 24 hours. Following dN treatment, 5×10^5^ HSPCs were electroporated with PE3 RNAs targeting *ATP1A1* and cultured in the presence or absence of 100 µM each dN and the indicated concentration of Vpx VLPs (GPP) or vehicle control for 72 hours. Genomic DNA was harvested 3 days post-nucleofection. *n* = 1 experiment. (**c**) PE and indels quantification as determined by BEAT and TIDE analysis from Sanger sequencing. 2.5×10^5^ HSPCs were electroporated with PE3max RNAs targeting *ATP1A1* and treated with 50 µM of the indicated dNs. Genomic DNA was harvested 3 days post-nucleofection. *n* = 3 independent biological replicates performed with CD34^+^ HSPCs from three different donors. (**d**) PE and indels quantification as determined by CRISPResso2 analysis from amplicon sequencing. 2.5×10^5^ HSPCs were electroporated with PE3/PE3bmax RNAs targeting *HBB* and cultured in the presence or absence of 5X Vpx VLPs (GPP) and 50 µM each dNs. Genomic DNA was harvested 3 days post-nucleofection. *n* = 3 independent biological replicates performed with CD34^+^ HSPCs from one donor homozygous for the rs713040 allele (donor 2). Each biological replicate is illustrated with the same shade of grey. (**e**) TwinPE and indel quantification as determined by CRISPResso2 analysis from amplicon sequencing. 5×10^5^ quiescent HSPCs were electroporated directly after thawing with Vpx mRNA and TwinPEmax RNAs targeting *AAVS1* and cultured in the presence or absence of 50 µM each dNs. Genomic DNA was harvested 3 days post-nucleofection. *n* = 3 independent biological replicates performed with CD34^+^ HSPCs from three different donors. Donor 2, circle. Donor 3, diamond. Donor 4, square. Donor 5, triangle. Donor 6, half-empty diamond. Donor 7, triangle (facing down).

The accessory lentiviral protein Vpx, encoded by HIV-2 and SIV viruses, associates with the CRL4^DCAF1^ E3 ubiquitin ligase to target SAMHD1 for proteasomal degradation^8^. We first tested whether delivering Vpx via virus-like particle (VLP) transduction could enhance prime editing. We cultured HSPCs in the presence of cytokines for 24 hours, and electroporated cells with a first-generation prime editor (PE2) mRNA and a previously optimized synthetic epegRNA/nick sgRNA pair targeting *ATP1A1*^11^. Following electroporation, we cultured HSPCs for 72 hours in the presence or absence of Vpx VLPs. The frequency of alleles harboring the *ATP1A1*-Q118R mutation increased from an average of 17% to 31% with HSPCs from two different healthy donors, as determined by Sanger sequencing (**Fig. 1b** and **Supplementary Fig. 1a**).

Since HSPCs lack the capacity to carry out de novo nucleotide synthesis and rely on membrane transporters such as ENT1^12^ to ensure extracellular nucleoside uptake and dNTP synthesis via the nucleoside salvage pathway, we hypothesized that exogenous deoxynucleoside (dN) supplementation could further boost prime editing (**Fig. 1a**). We treated HSPCs with 100 µM of each dN before and after electroporation and observed an increase in prime editing efficiency from 15% to 42% without Vpx VLPs, and to 48% with Vpx VLPs (**Fig. 1b**). We then treated cells with different concentrations of dNs before, after, and both before and after electroporation. Although dNs enhanced PE with all conditions, higher improvements (from 18% to 42%) were observed when supplementing dNs after electroporation (**Supplementary Fig. 1b**). For subsequent experiments, cells were treated with or without 50 µM of each dN after electroporation to maximize prime editing efficiency.

We then tested whether all four dNs were needed for maximal PE enhancement. To further boost efficiency, we generated PEmax^2^ mRNA which markedly increased the basal level of editing to an average of 41% *ATP1A1*-Q118R allele frequency across HSPCs from three different donors (**Fig. 1c**). We observed an increase with all individual dNs as well as all dNs minus one conditions, but the highest level of editing was observed with all four dNs to an average of 64% edited alleles (**Fig. 1c**). Since all four dNTPs are needed to synthesize the *ATP1A1*-Q118R epegRNA flap, these results are consistent with the hypothesis that the dNTP concentration is limiting for efficient reverse transcription and PE in HSPCs. We also tested the impact of dNs supplementation and overexpression of Vpx, SAMHD1, or the inactive SAMHD1-H215A variant on PE in K562 and Jurkat cells. Modulating the nucleotide metabolism had little to no impact on editing efficiency, suggesting that the concentration of dNTPs is not limiting in cancer cell lines (**Supplementary Fig. 2**).

We then assessed the impact of dN supplementation and Vpx VLPs using an additional epegRNA designed to correct the *HBB*-E6V mutation causative of sickle cell disease using PE3 and PE3b strategies^13^. While Vpx VLP and dN treatments improved PE at *ATP1A1* (**Supplementary Fig. 3**), modest improvements were observed at the *HBB* locus with the PE3 strategy (**Fig. 1d**). Nonetheless, the average level of editing increased from 35% to 42% with an average of just 1.25% indels with the PE3b strategy, as determined by amplicon sequencing (**Fig. 1d**). This level of product purity contrasted with the high level of tandem duplications observed at *ATP1A1* with the PE3 strategy in the presence or absence of dNs and Vpx VLPs (**Supplementary Fig. 3**), corroborating previous findings in K562 cells with pegRNA and nick sgRNA PAMs facing outward towards each other (PAM-out configuration)^11^.

Editing quiescent HSCs without cytokine culture reduces DSB genotoxicity^14^ and may ultimately enable editing HSCs *in vivo*. We hypothesized that the nucleotide pool available for reverse transcription could especially limit PE in quiescent HSPCs that had not been stimulated by cytokines *in vitro*. We also generated Vpx mRNA to co-deliver at an equimolar ratio with PEmax mRNA during electroporation to confirm that Vpx alone could ameliorate PE (**Supplementary Fig. 4a**). We performed electroporation directly after thawing CD34+ HSPCs, and cultured cells for 72 hours in the presence or absence of 50 µM dNs (**Supplementary Fig. 4a**). While Vpx mRNA co-delivery and dN supplementation improved editing at *ATP1A1*, modulation of nucleotide metabolism had minimal impact on PE in quiescent HSPCs with the *HBB* epegRNA and an additional epegRNA designed to knockout *B2M*^15^ (**Supplementary Fig. 4b**,**c**). Of relevance, we noted that the highest level of improvement was observed with the *ATP1A1*-Q118R_v2 epegRNA, which was optimized to evade DNA mismatch repair (MMR)^11^ while the *HBB* and *B2M* epegRNAs introduce a single nucleotide substitution and are thus sensitive to MMR. Since MMR antagonizes prime editing efficiency and fidelity^2,16^, we hypothesized that modulating the nucleotide metabolism could positively interact with MMR evasion to facilitate prime editing in HSPCs.

Twin prime editing (TwinPE) generates 3’ DNA flaps dissimilar to the target site and is not expected to produce heteroduplexes that engage MMR. We next tested the impact of modulating nucleotide metabolism on TwinPE using a pair of epegRNAs designed to install a Bxb1 *attP* recombinase site at the *AAVS1* locus^3^. Vpx mRNA co-delivery and dN supplementation markedly improved TwinPE efficiency at *AAVS1* in quiescent HSPCs (**Fig. 1e**). Despite the considerable variability observed between quiescent HSPCs from different donors, modulating the nucleotide metabolism improved TwinPE consistently, with a 2.3 to 4.2-fold increase across three donors. Consistent with the hypothesis of dNTPs being especially limiting in quiescent HSPCs, a higher basal level of twin prime editing and a reduced improvement with Vpx mRNA and dN supplementation were observed in HSPCs cultured with cytokines before electroporation (**Supplementary Fig. 5**). Since TwinPE bypasses multiple DNA repair steps that can reject traditional prime edits, including 3’ flap annealing to the genome and heteroduplex resolution^3,17,18^, these results suggest that MMR evasion interacts positively with dN supplementation and Vpx to boost prime editing efficiency.

Considering the lower product purity observed when using an additional nick sgRNA, we next tested whether modulating the nucleotide metabolism could positively interact with MMR evasion to achieve high levels of editing with the PE2 approach. A simple yet very efficient approach to evade MMR is to install additional silent mutations near the intended edit^2,19^, as previously demonstrated with *ATP1A1*-Q118R v1-v3 epegRNAs^11^ (**Fig. 2a**). In HSPCs, we observed a marked increase in PE2 efficiency with three (v2) and four (v3) substitutions compared to two (v1) using *ATP1A1*-Q118R_v1-v3 epegRNAs (**Fig. 2a**), corroborating previous findings in K562 cells^11^. Treatment with Vpx VLPs and dNs further enhanced precise prime editing from an average of 20.1% to 42.9% allele frequency with the v3 epegRNA, allowing up to 53.2% PE alleles with 0.4% indels and completely abrogating tandem duplications, as determined by amplicon sequencing, i.e. >100 PE-to-indel purity ratio (**Fig. 2a** and **Supplementary Fig. 6**). We then designed additional *B2M* epegRNAs introducing C•C mismatches which are resistant to MMR^2,20–22^ (**Fig. 2b**). Strikingly, PE efficiency rose from undetectable levels with the original *B2M* epegRNA^15^ to an average of 25% and 61% with one and two C•C mismatches, respectively (**Fig. 2b**). Co-delivering Vpx mRNA and supplementing dNs steadily increased PE with all MMR-evading epegRNAs, although to a lower extent than the *ATP1A1*-Q118R_v3 epegRNA (**Fig. 2a,b**). Together, MMR evasion and nucleotide metabolism modulation allowed up to 71% prime editing at *B2M* (**Fig. 2b**). These data demonstrate that MMR opposes prime editing in HSPCs and that modulating nucleotide metabolism positively interacts with MMR-evasion to install prime edits with high purity and efficiency in the absence of additional nicking sgRNAs using the PE2max system.

**Figure 2.**
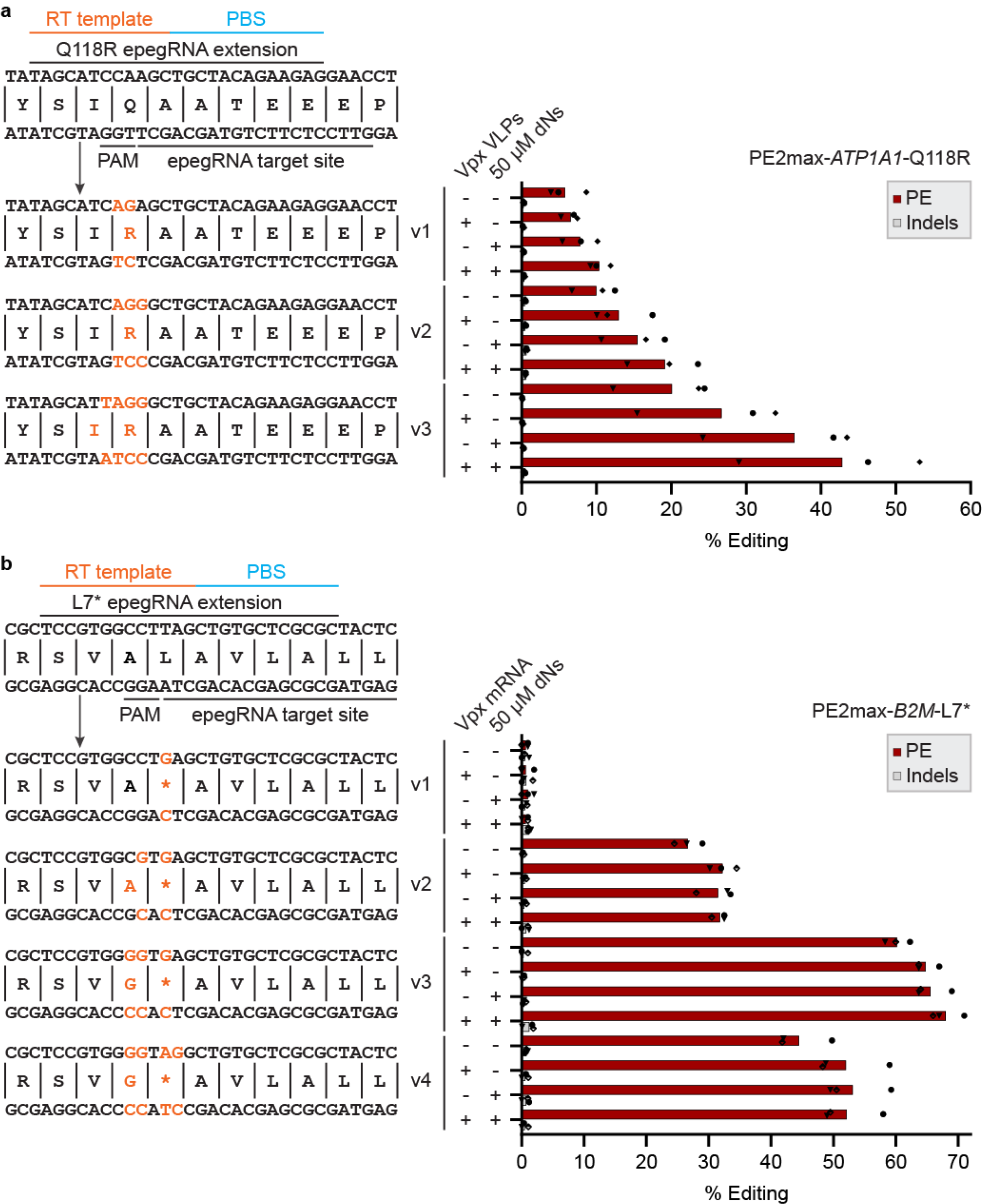
Modulation of nucleotide metabolism positively interacts with MMR evasion. (**a**) PE and indel quantification as determined by CRISPResso2 analysis from amplicon sequencing. The MMR evasion strategy based on the installation of additional silent mutations and the epegRNA used to target the *ATP1A1* locus are illustrated on the left. Point mutations are illustrated in orange. 2.5×10^5^ HSPCs were electroporated with PE2max RNAs targeting *ATP1A1* and cultured in the presence or absence of 50 µM each dN and 25X in-house Vpx VLPs. Genomic DNA was harvested 3 days post-nucleofection. *n* = 3 independent biological replicates performed with CD34^+^ HSPCs from three different donors. (**b**) PE and indels quantification as determined by BEAT and TIDE analysis from Sanger sequencing. 2.5×10^5^ HSPCs were electroporated with Vpx mRNA and PE2max RNAs targeting *B2M* and cultured in the presence or absence of 50 µM each dN. Genomic DNA was harvested 3 days post-nucleofection. *n* = 3 independent biological replicates performed with CD34^+^ HSPCs from three different donors. Donor 2, circle. Donor 6, half-empty diamond. Donor 7, triangle (facing down).

The enhancements observed in this study suggest that the low concentration of dNTPs available for reverse transcription restricts prime editing in HSPCs. The marked positive interactions between the modulation of nucleotide metabolism and TwinPE could be interpreted that boosting the level of dNTPs may facilitate both reverse transcription and the DNA polymerase fill-in synthesis step required to install TwinPE modifications. Paradoxically, dNTP availability could also boost MMR, which depends on fill-in synthesis by DNA polymerase δ after the excision of the mismatched DNA strand^18,23,24^, and antagonize PE by restoring the original sequence more efficiently. This could partially explain the greater increase in PE efficiency observed with modulating nucleotide metabolism along with MMR-evading mutations or twin prime editing.

Altogether, modulation of nucleotide metabolism enhances prime editing in HSPCs which should facilitate clinical translation of therapeutic prime editing. Combining modulation of nucleotide metabolism, PEmax editors, and MMR evasion designs allows unprecedented prime editing efficiency and purity in HSPCs. Considering recent progress towards delivery of genome editors to HSCs *in vivo*^25,26^, these findings may help designing novel prime editing therapies to engineer quiescent HSCs *in situ*.

## METHODS

### Cell culture

Cryopreserved human CD34^+^ HSPCs from mobilized peripheral blood of deidentified healthy donors were obtained from the Fred Hutchinson Cancer Research Center (Seattle, Washington). CD34^+^ HSPCs were cultured with X-Vivo-15 media supplemented with 100 ng/ml human Stem Cell Growth Factor (SCF), 100 ng/ml human thrombopoietin (TPO), and 100 ng/ml recombinant human FMS-like Tyrosine Kinase 3 Ligand (Flt3-L). K562 (CCL-243) and Jurkat (TIB-152) cells were obtained from the ATCC and cultured at 37^°^C under 5% CO2 in RPMI media supplemented with 10% FBS, and Penicillin/Streptomycin. HEK293T (CRL-1573) cells were obtained from ATCC and cultured at 37^°^C under 5% CO2 in DMEM media supplemented with 10% FBS, and Penicillin/Streptomycin. Deoxynucleosides (Sigma-Aldrich) were resuspended in water at 12.5 mM each, filter-sterilized, and stored at -20^°^C.

### Vpx virus-like particles (VLPs)

Vpx VLPs were provided by the Genetic Perturbation Platform (GPP) of the Broad Institute of MIT and Harvard or produced in-house. Briefly, HEK293T cells were transfected with VSV-G envelope and SIV Vpx vectors, and virus-like particles were harvested from the supernatant 35-40 hours post-lipofection. The 1X concentration (1/20 of the culture volume) was established as the minimal volume to enhance lentiviral transduction of primary human monocytes^27^. Titration of activity was performed using prime editing and cell viability as readouts, and a concentration of 5X (1/4 of the culture volume) was used for all experiments using GPP VLPs. HEK293T culture media was used as a vehicle control in all experiments using GPP VLPs. Alternatively, in-house VLPs were produced using the same protocol followed by ultracentrifugation at 24,000 RPM for 2 hours (SW 28 Beckman swinging bucket rotor) to concentrate the VLPs and avoid diluting the HSPCs culture media. Titration of activity was performed using prime editing and cell viability as readouts, and a concentration of 25X Vpx VLPs (normalized from the initial volume of producer cell line supernatant) was used for all experiments using in-house VLPs.

### *In vitro* transcription, and epegRNA and nick sgRNA synthesis

The PE2 and PEmax transcription template plasmids were linearized, and mRNA was transcribed using the HiScribe T7 high yield RNA kit (NEB) using N1-methylpseudouridine (Trilink) instead of uridine, and co-transcriptional capping with CleanCap AG (Trilink). For Vpx mRNA IVT template, a cassette harboring a 5’ UTR region with an eIF4G aptamer, a kozak sequence, the Vpx cassette, and a 3’ UTR harboring the WPRE element was cloned in the pT7-PEmax for IVT plasmid (Addgene 178113)^2^. The template was generated as previously described^2^. Briefly, the template was PCR amplified with a forward primer that correct a T7 promoter inactivating mutation and a reverse primer that appends a 119-nt poly(A) tail to the 3’ UTR. Following IVT, mRNAs were purified using the Monarch RNA Cleanup kit (500 µg) (NEB) and eluted in 1X nuclease-free IDTE buffer (10 mM Tris, 0.1 mM EDTA, pH 7.5). The mRNA concentration was quantified using Qubit RNA high sensitivity (HS) kit (ThermoFisher). Synthetic epegRNAs were provided by Integrated DNA Technologies (IDT) and resuspended at 200 pmol/µl in nuclease-free IDTE buffer (10 mM Tris, 0.1 mM EDTA, pH 7.5). The epegRNA contained 2’-O-methyl modifications at the three first and the last three nucleotides, and phosphorothioate linkages between the three first and the last three nucleotides. A UUU trinucleotide was appended to the 3’ end of each epegRNA after the tevopreQ1 motif. Synthetic nick sgRNAs were provided by IDT at a 10 nmol scale and resuspended at 200 pmol/µl in nuclease-free IDTE buffer (10 mM Tris, 0.1 mM EDTA, pH 7.5).

### Genome editing vectors

Prime editing experiments in K562 and Jurkat cell lines were performed with pCMV-PEmax (Addgene 174820). The *ATP1A1*-Q118R_v2 epegRNA^11^ was cloned into pU6-tevopreq1-GG-acceptor (Addgene 174038) and the *ATP1A1*-G3 nick sgRNA^11^ was cloned into SpCas9_sgRNA_expression_in_pBluescript (Addgene 122089). For SIV-Vpx, SAMHD1, and the inactive SAMHD1-H215A variant overexpression under the transcriptional control of the CMV promoter, the PEmax cassette from pCMV-PEmax (Addgene 174820) was swapped with the indicated cassette.

### Nucleofection

For standard conditions, CD34^+^ HSPCs were thawed and cultured for 24 hours in the presence of cytokines prior to nucleofection. CD34^+^ HSPCs were electroporated using the P3 Primary Cell X kit S (Lonza) according to manufacturer’s recommendations. Unless otherwise indicated, 2.5×10^5^ cells were electroporated with 2000 ng PE mRNA, 200 pmol epegRNA, and 100 pmol nick sgRNA using pulse code DS-130. For twin prime editing conditions, 2.5×10^5^ cells were electroporated with 2000 ng PE mRNA and 150 pmol of each epegRNA using pulse code DS-130. For Vpx mRNA co-delivery, Vpx:PEmax mRNAs were electroporated at an equimolar ratio. For quiescent HSPCs nucleofections, HSPCs where thawed and 5×10^5^ cells were electroporated without prior culture with cytokines. Following electroporation, 80 µl of media supplemented with cytokines was added to each well and cells were incubated for 10 minutes prior to transfer to the culture plate. Were indicated, cells from one nucleofection were split in two wells in a 96-well plate with or without the indicated concentration of each deoxynucleosides. For Vpx VLPs experiments, the culture media was supplemented with 1X Lentiboost®. DMEM supplemented with 10% FBS and 1% Penicillin/Streptomycin was used as a vehicle control for Vpx VLPs from the GPP platform. CD34^+^ HSPCs were cultured for 72 hours before genotyping.

K562 cells (2×10^5^ cells) were electroporated with 750 ng PEmax vector, 250 ng epegRNA vector, and 100 ng nick sgRNA vector using the SF nucleofection kit (Lonza) and pulse code FF-120. Jurkat cells (5×10^5^ cells) were electroporated with 500 ng PEmax vector, 250 ng epegRNA vector, and 100 ng nick sgRNA vector using the SE nucleofection kit (Lonza) and pulse code CL-120. K562 and Jurkat cells were cultured for 72 hours before genotyping.

### Genotyping

Three days post-nucleofection, cells were washed once with 500 µl PBS and genomic DNA was harvested using QuickExtract DNA extraction solution (Epicentre) following manufacturer’s recommendations. For Sanger sequencing, PCR amplifications were performed with 30 cycles of amplification with Phusion high-fidelity polymerase. Primers used in this study and the amplicon sizes are provided in the Supplementary material section. For amplicon sequencing, PCR amplifications were performed with Phusion high-fidelity polymerase or KOD Hot Start DNA Polymerase and PCR product quality was evaluated by electrophoresis. One µl of locus-specific PCR product was used for indexing (PCR 2) with KOD Hot Start DNA polymerase. Amplicon PCR products were purified using AMPure magnetic beads, and quality was evaluated by electrophoresis and TapeStation using D1000 high sensitivity Screen tape (Agilent). Amplicons were sequenced using paired-end 150 bp reads on an Illumina NovaSeq X system by Novogene (Durham, NC, USA) or in-house on an Illumina MiniSeq system. The percentage of prime edited alleles and indels were quantified using BEAT^28^ and TIDE^29^ webtools from Sanger sequence data files or CRISPResso2^30^ from amplicon sequencing data files. The percentage of indel reads was determined as the percentage of modified reads and substitutions were excluded for the analysis. For twin prime editing, CRISPResso2 was run in HDR mode using the desired allele as the expected allele and the percentage of indels was determined as the % NHEJ reads + % Imperfect HDR reads, and substitutions were excluded for indels quantification.

## DATA AVAILABILITY

All sequencing data will be publicly accessible from the National Center for Biotechnology Information Bioproject database and the accession number will be available before publication. Vectors used in this study will be publicly available via Addgene before publication.

## ACKNOWLEDGEMENTS

This work was supported by the Doris Duke Foundation. CD34^+^ cells were provided by Fred Hutch Cooperative Center of Excellence in Hematology. We thank Denise Klatt and Christian Brendel for their support in producing in-house Vpx virus-like particles, the Genetic Perturbation Platform (GPP) of the Broad Institute of MIT and Harvard for providing VSV-G envelope and SIV Vpx vectors, and Andrea Consentino and Pietro Genovese for generously providing the PEmax vector used for *in vitro* transcription.

## AUTHOR CONTRIBUTIONS

Conceptualization, S.L. and D.E.B.; Methodology, S.L. and D.E.B.; Investigation, S.L. A.V., and D.E.B.; Original Draft, S.L.; Writing, Review, and Editing, S.L. and D.E.B.; Funding Acquisition, D.E.B.; Supervision, D.E.B.

## COMPETING INTERESTS

S.L. and D.E.B. have filed a provisional patent application related to this work.

**Supplementary Figure 1.**
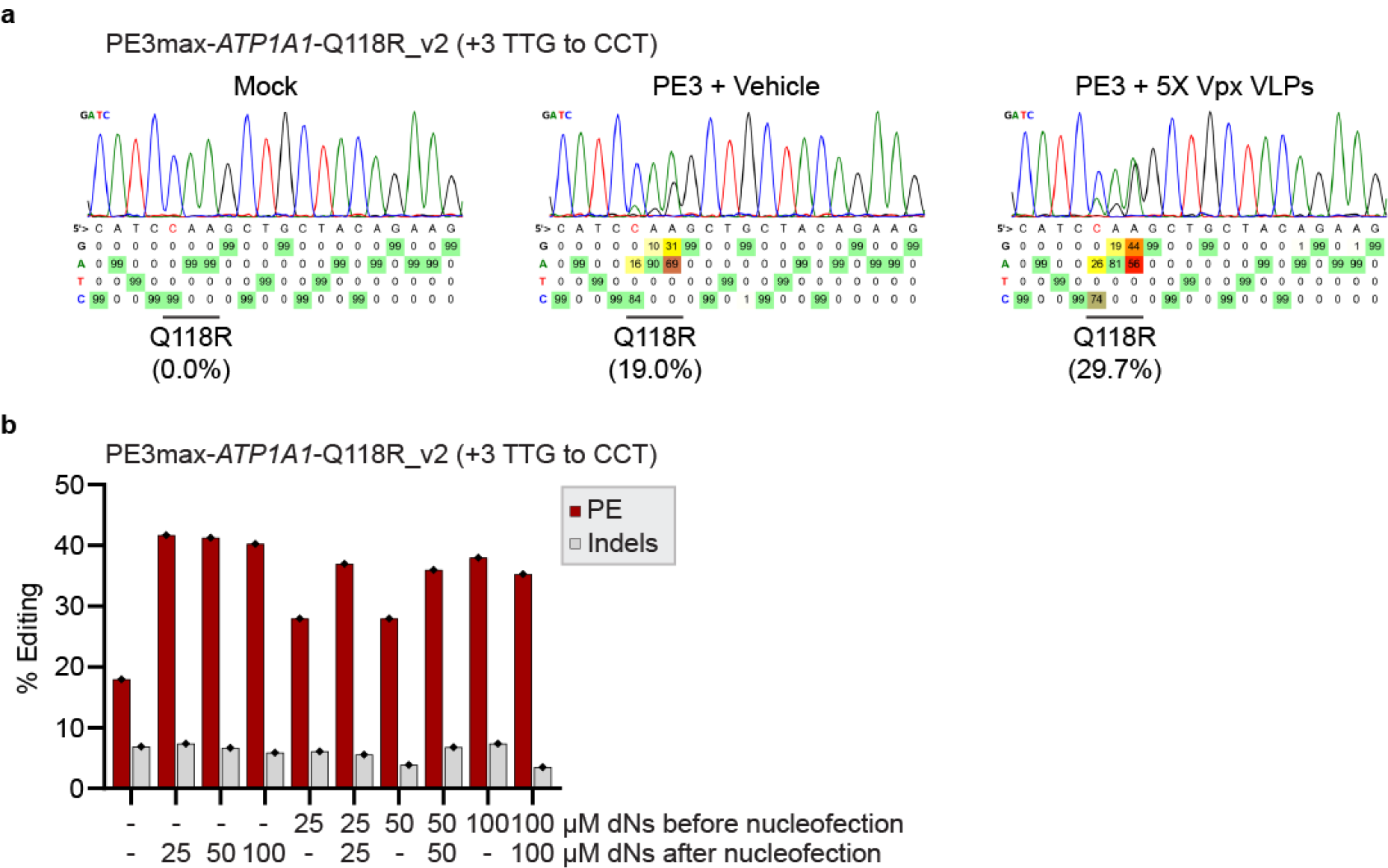
Vpx VLP and dN supplementation enhance prime editing in HSPCs with the first-generation PE2 prime editor. (**a**) PE quantification as determined by BEAT analysis from Sanger sequencing. 5×10^5^ HSPCs were electroporated with PE3 RNAs targeting *ATP1A1* and treated with 5X Vpx VLP (GPP) or vehicle control. Genomic DNA was harvested 3 days post-nucleofection. *n* = 1 experiment (donor 1) replicated with HSPCs from donor 2 with similar results (See **Fig. 1b**). (**b**) PE and indels quantification as determined by BEAT and TIDE analysis from Sanger sequencing. HSPCs were thawed and cultured in the presence or absence of the indicated concentration of each dN for 24 hours. Following dN treatment, 1.25×10^5^ HSPCs were electroporated with PE3 RNAs targeting *ATP1A1* and cultured in the presence or absence of the indicated concentration of each dN for 72 hours. Genomic DNA was harvested 3 days post-nucleofection. *n* = 1 experiment. Donor 3, diamond.

**Supplementary Figure 2.**
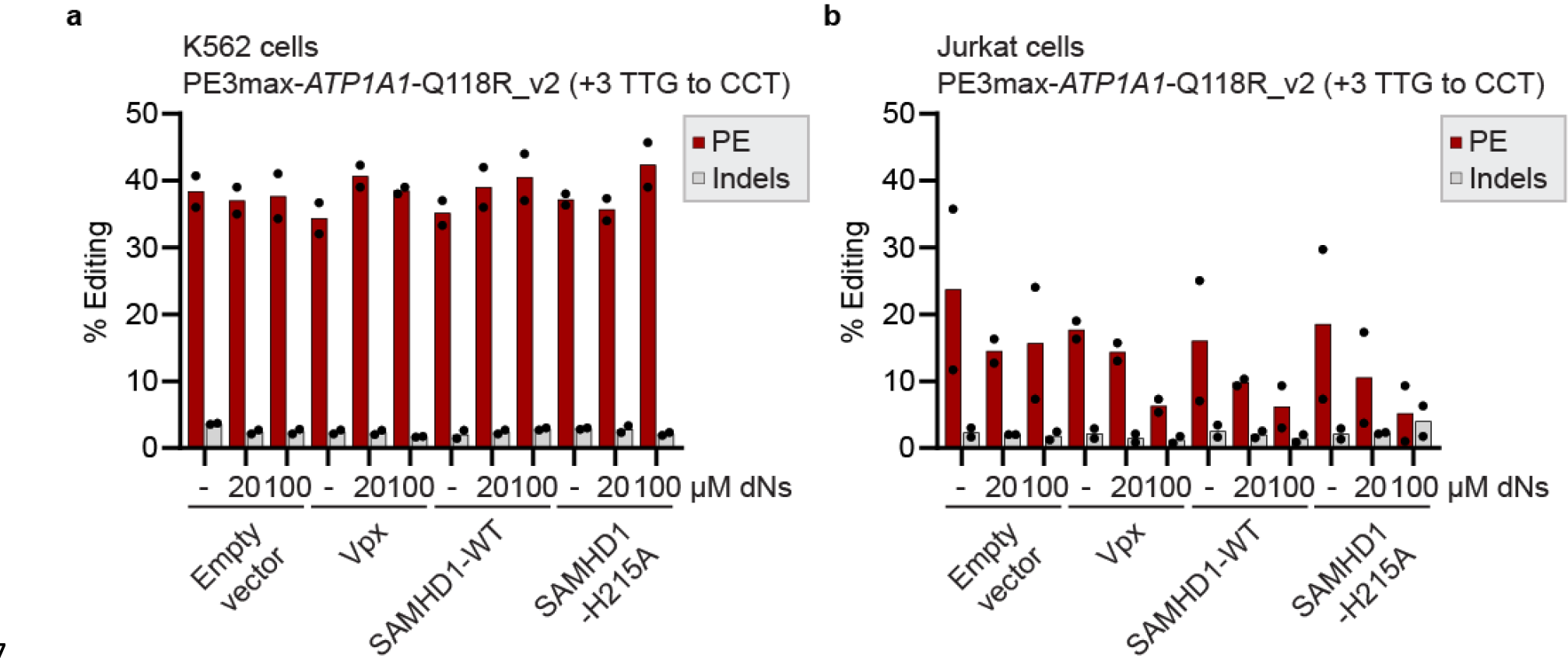
Modulation of nucleotide metabolism has minimal impact on prime editing in cancer cell lines. (**a**) PE and indels quantification as determined by BEAT analysis from Sanger sequencing. K562 cells were electroporated with PE3max vectors targeting *ATP1A1* and the indicated Vpx or SAMHD1 vector and cultured with the indicated concentration of each dN for 72 hours. An empty pUC19 vector was used as a negative control to normalize the concentration of DNA in all nucleofections. Genomic DNA was harvested 3 days post-nucleofection. *n* = 2 independent biological replicates performed at different times with equivalent results. (**b**) Same as in (**a**) with Jurkat cells.

**Supplementary Figure 3.**
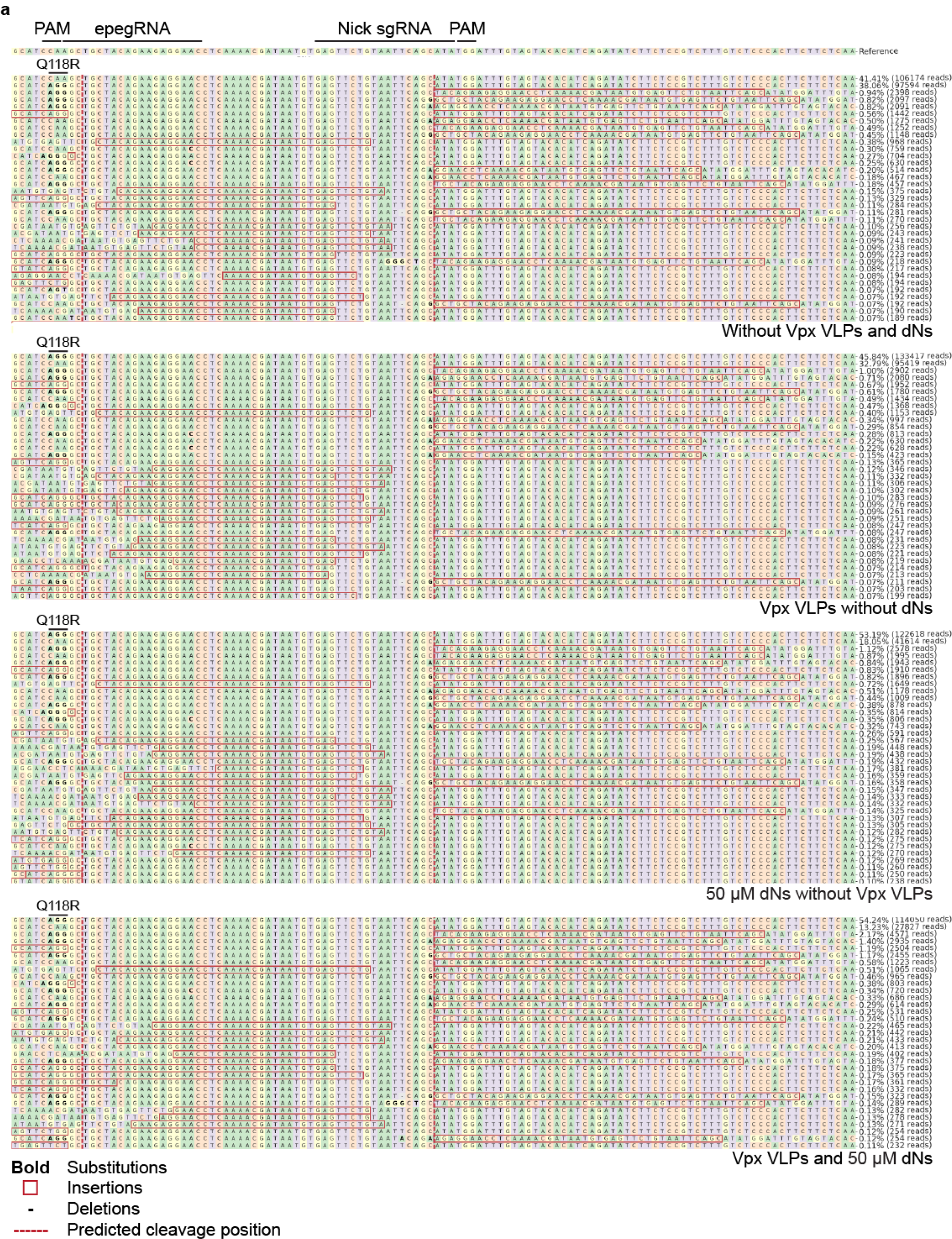
Staggered PE3 nicks generate tandem duplications at *ATP1A1* in HSPCs. CRISPResso2 amplicon sequencing allele plots after PE3 at *ATP1A1*. 2.5×10^5^ HSPCs were electroporated with PE3max RNAs targeting *ATP1A1* (Q118R_v2, +3 TTG to CCT) and cultured in the presence or absence of 5X Vpx VLPs (GPP) and 50 µM each dN. Genomic DNA was harvested 3 days post-nucleofection. Representative allele plots are from one of three independent biological replicates performed with CD34^+^ HSPCs from three different donors with equivalent results. Insertions are illustrated in red squares, and they represent tandem duplications of the genomic sequence found between the two nick sites.

**Supplementary Figure 4.**
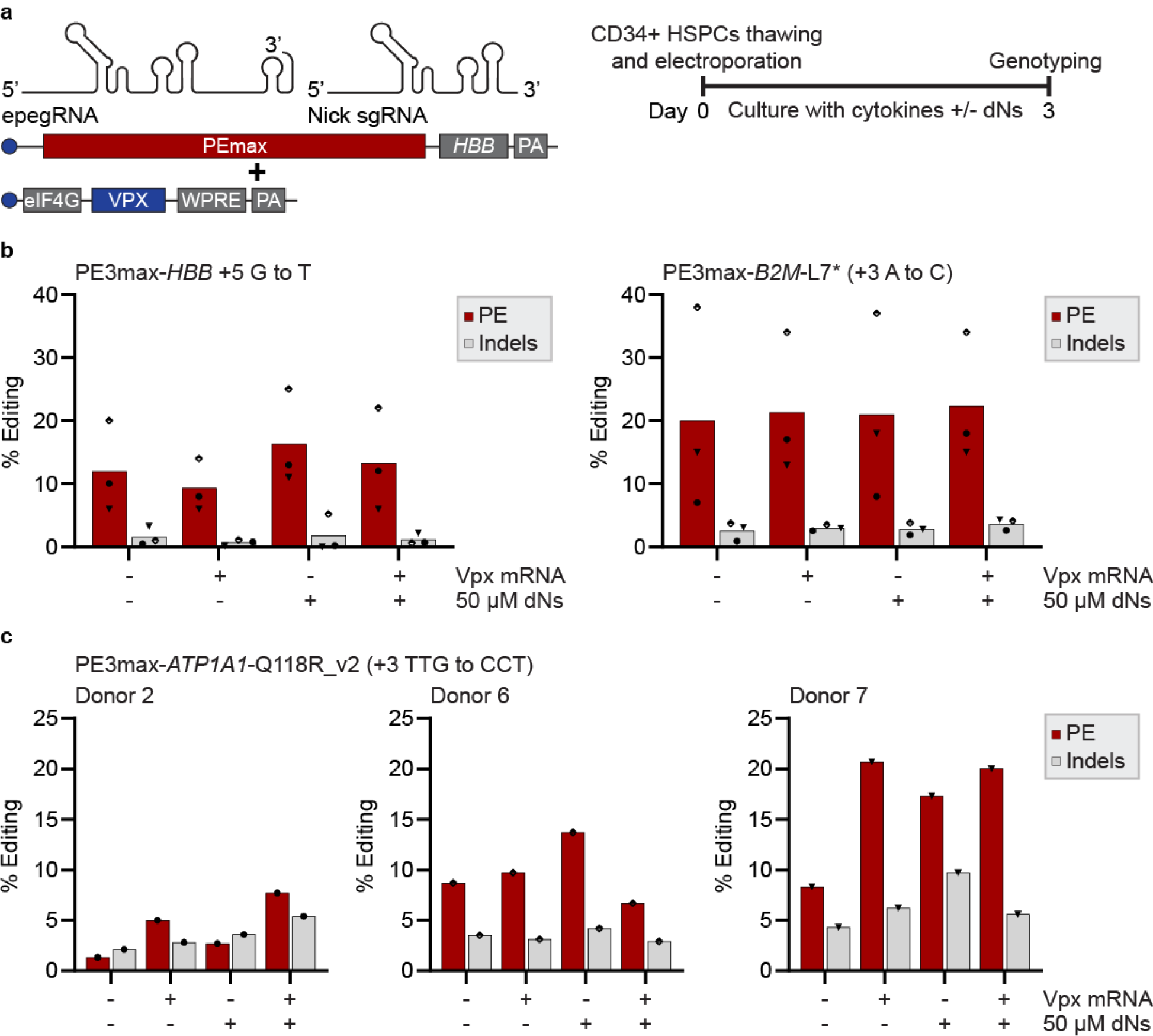
Modulation of nucleotide metabolism positively interacts with MMR-evading epegRNAs in quiescent HSPCs. (**a**) Schematic representation of the RNAs used to co-deliver Vpx with PE3max components. The timeline for prime editing in quiescent CD34^+^ HSPCs is illustrated on the right. (**b**) PE and indels quantification as determined by BEAT and TIDE analysis from Sanger sequencing. 5×10^5^ quiescent HSPCs were electroporated directly after thawing with Vpx mRNA and PE3max RNAs targeting *HBB* or *B2M* and cultured in the presence or absence of 50 µM each dN. Genomic DNA was harvested 3 days post-nucleofection. *n* = 3 independent biological replicates performed with CD34^+^ HSPCs from three different donors. (**c**) Same as in (**b**) for prime editing at *ATP1A1* with MMR-evading substitutions. Donor 2, circle. Donor 6, half-empty diamond. Donor 7, triangle (facing down).

**Supplementary Figure 5.**
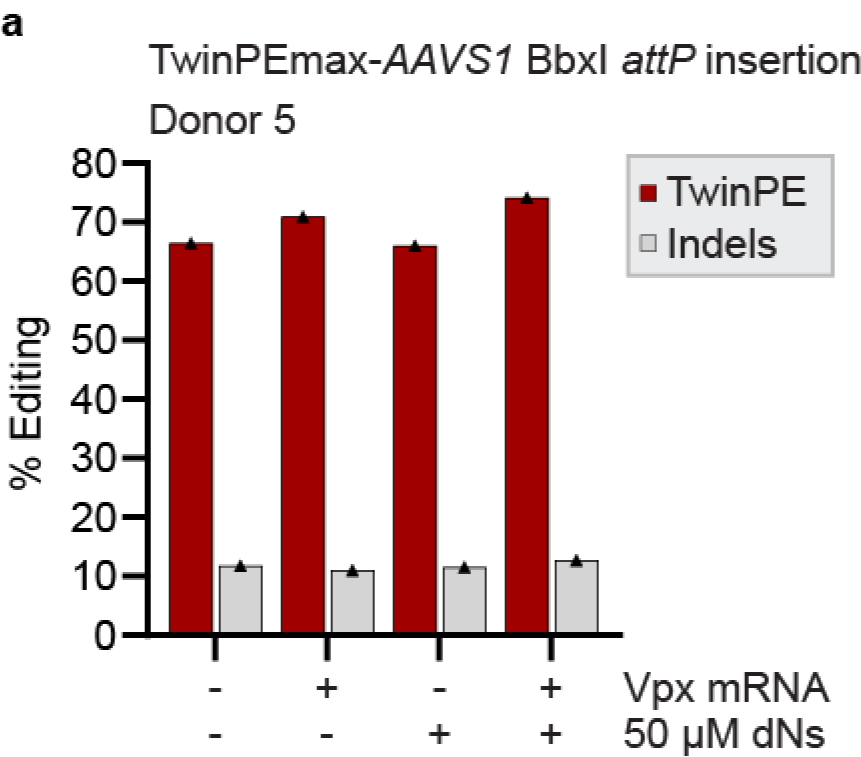
*Ex vivo* HSPCs culture with cytokines moderates the impact of Vpx and dNs supplementation on twin prime editing efficiency. (**a**) TwinPE and indel quantification as determined by CRISPResso2 analysis from amplicon sequencing. After 24 hours of culture with cytokines, 2.5×10^5^ quiescent HSPCs were electroporated with Vpx mRNA and TwinPEmax RNAs targeting *AAVS1* and cultured in the presence or absence of 50 µM each dNs. Genomic DNA was harvested 3 days post-nucleofection. *n* = 1 experiment. Donor 5, triangle.

**Supplementary Figure 6.**
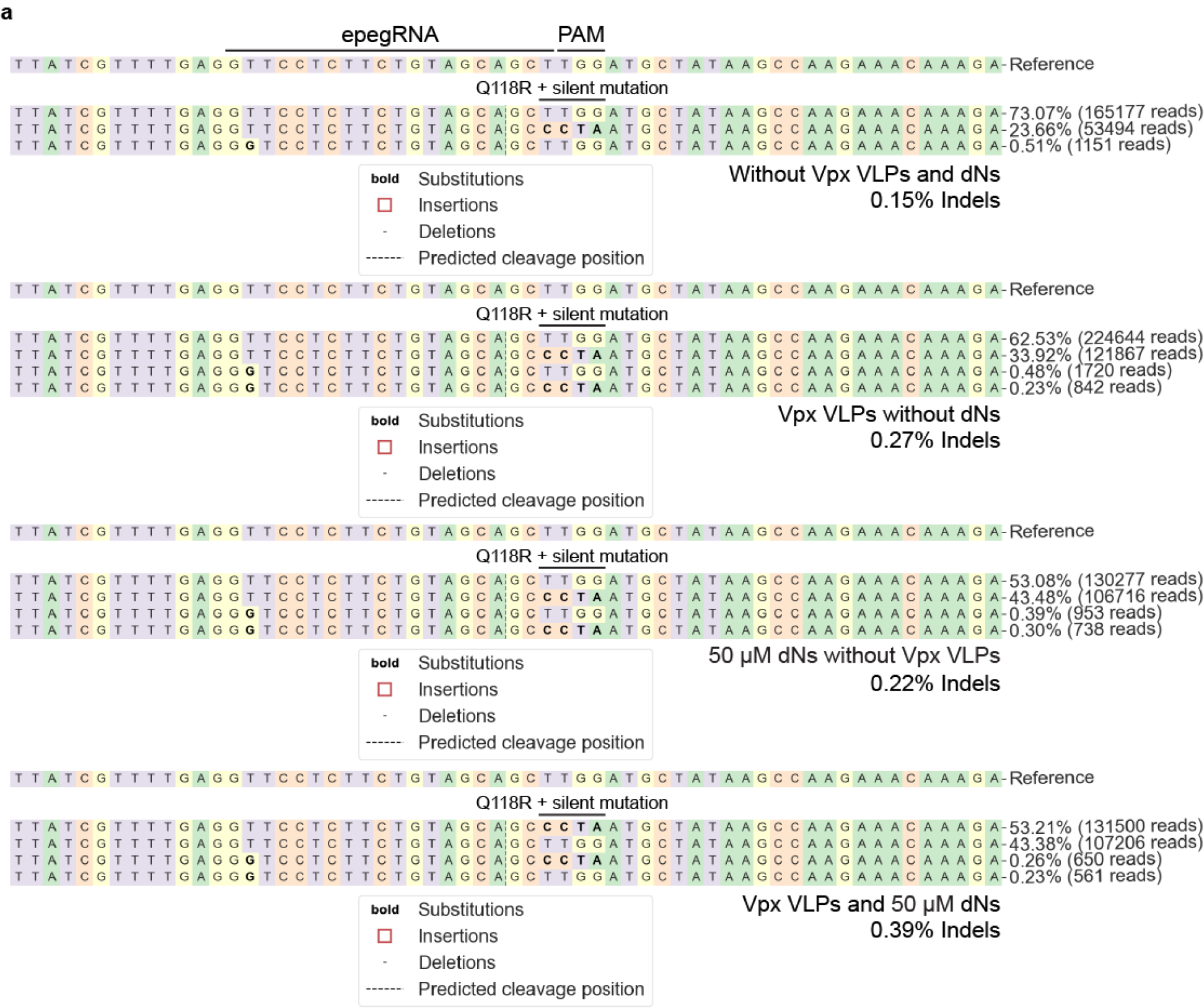
Omitting the nick sgRNA abrogates indels and tandem duplications at *ATP1A1* in HSPCs. (**a**) CRISPResso2 amplicon sequencing allele plots from **Fig. 2a**. 2.5×10^5^ HSPCs were electroporated with PE2max RNAs targeting *ATP1A1* (Q118R_v3, +3 TTGG to CCTA) and cultured in the presence or absence of 25X in-house Vpx VLPs and 50 µM each dN. Genomic DNA was harvested 3 days post-nucleofection. Representative allele plots are from one of three independent biological replicates performed with CD34^+^ HSPCs from three different donors with equivalent results. Reads with ≥ 0.2% frequency are shown, and the frequency of indels is indicated for each sample.

