## Supplemental information for "Nucleotide metabolism constrains prime editing in hematopoietic stem and progenitor cells"

### Supplementary information - Levesque, Verma, and Bauer 2023

**Primers used in this study for Sanger sequencing**

| Primer | Sequence | PCR product size |
|---|---|---|
| *ATP1A1* -Forward | TCTGGGTGTTATGAGTTCCTTGGG | 740 bp |
| *ATP1A1* -Reverse | AACATGAATCGCTGTGCTCTGC | |
| *HBB* -Forward | CTGAGGGTTTGAAGTCCAACTCC | 760 bp |
| *HBB* -Reverse | CCTGTTACTTATCCCCTTCCTATGAC | |
| *AAVS1* -Forward | CAGCTAGTCTTCTTCCTCCAACC | 743 bp |
| *AAVS1* -Reverse | GACGCACCATTCTCACAAAGG | |
| *B2M* -Forward | CCTCCAGCCTGAAGTCCTAGAATG | 713 bp |
| *B2M* -Reverse | GCGAAAGAGCGGAAGAGAAACC | |

**Primers used in this study for Sanger sequencing**

| Primer | Sequence | PCR product size |
|---|---|---|
| *ATP1A1* -NGS-Forward | CTACACGACGCTCTTCCGATCTGGGGGTTCTCAATGTTACTGTGG | 281 bp |
| *ATP1A1* -NGS-Reverse | AGACGTGTGCTCTTCCGATCTGAAGGCAACAGCTGTTCATAACC | |
| *HBB* -NGS-Forward | CTACACGACGCTCTTCCGATCTAGGGCTGGGCATAAAAGTCAG | 278 bp |
| *HBB* -NGS-Reverse | AGACGTGTGCTCTTCCGATCTTGCCCAGTTTCTATTGGTCTCC | |
| *AAVS1* -NGS-Forward | CTACACGACGCTCTTCCGATCTCATTTCCTGGAGCCATCTCTCTC | 283 bp |
| *AAVS1* -NGS-Reverse | AGACGTGTGCTCTTCCGATCTTCTTGGGAAGTGTAAGGAAGCTG | |

**epegRNAs and nick sgRNAs used in this study**

| epegRNA | Sequence |
|---|---|
| *ATP1A1* _Q118R_v1 | GUUCCUCUUCUGUAGCAGCUGUUUUAGAGCUAGAAAUAGCAAGUUAAAAUAAGGCUAGUCCGUUAUCAACUUGAAAAAGUGGCACCGAGUCGGUGCUAGCAUCAGAGCUGCUACAGAAGAGCGCGGUUCUAUCUAGUUACGCGUUAAACCAACUAGAAUUU |
| *ATP1A1* _Q118R_v2 | GUUCCUCUUCUGUAGCAGCUGUUUUAGAGCUAGAAAUAGCAAGUUAAAAUAAGGCUAGUCCGUUAUCAACUUGAAAAAGUGGCACCGAGUCGGUGCUAGCAUCAGGGCUGCUACAGAAGAGCGCGGUUCUAUCUAGUUACGCGUUAAACCAACUAGAAUUU |
| *ATP1A1* _Q118R_v3 | GUUCCUCUUCUGUAGCAGCUGUUUUAGAGCUAGAAAUAGCAAGUUAAAAUAAGGCUAGUCCGUUAUCAACUUGAAAAAGUGGCACCGAGUCGGUGCUAGCAUUAGGGCUGCUACAGAAGAGCGCGGUUCUAUCUAGUUACGCGUUAAACCAACUAGAAUUU |
| *HBB* _E6V_Correction | CAUGGUGCACCUGACUCCUGGUUUUAGAGCUAGAAAUAGCAAGUUAAAAUAAGGCUAGUCCGUUAUCAACUUGAAAAAGUGGCACCGAGUCGGUGCAGACUUCUCUUCAGGAGUCAGGUGCACAGAAUAAACGCGGUUCUAUCUAGUUACGCGUUAAACCAACUAGAAUUU |
| *AAVS1* _A1615a_TwinPE | GCAGCUCAGGUUCUGGGAGAGUUUUAGAGCUAGAAAUAGCAAGUUAAAAUAAGGCUAGUCCGUUAUCAACUUGAAAAAGUGGCACCGAGUCGGUGCUACCGUACACCACUGAGACCGCGGUGGUUGACCAGACAAACCUCCCAGAACCCGCGGUUCUAUCUAGUUACGCGUUAAACCAACUAGAAUUU |
| *AAVS1* _B1705b_TwinPE | GAUGGAGCCAGAGAGGAUCCGUUUUAGAGCUAGAAAUAGCAAGUUAAAAUAAGGCUAGUCCGUUAUCAACUUGAAAAAGUGGCACCGAGUCGGUGCGUCUGGUCAACCACCGCGGUCUCAGUGGUGUACGGUACAAACCUUCCUCUCUGGCCGCGGUUCUAUCUAGUUACGCGUUAAACCAACUAGAAUUU |
| *B2M* _L7Stop_v1 | GAGUAGCGCGAGCACAGCUAGUUUUAGAGCUAGAAAUAGCAAGUUAAAAUAAGGCUAGUCCGUUAUCAACUUGAAAAAGUGGCACCGAGUCGGUGCUCCGUGGCCUGAGCUGUGCUCGCGCCGCGGUUCUAUCUAGUUACGCGUUAAACCAACUAGAAUUU |
| *B2M* _L7Stop_v2 | GAGUAGCGCGAGCACAGCUAGUUUUAGAGCUAGAAAUAGCAAGUUAAAAUAAGGCUAGUCCGUUAUCAACUUGAAAAAGUGGCACCGAGUCGGUGCUCCGUGGCGUGAGCUGUGCUCGCGCCGCGGUUCUAUCUAGUUACGCGUUAAACCAACUAGAAUUU |
| *B2M* _L7Stop_v3 | GAGUAGCGCGAGCACAGCUAGUUUUAGAGCUAGAAAUAGCAAGUUAAAAUAAGGCUAGUCCGUUAUCAACUUGAAAAAGUGGCACCGAGUCGGUGCUCCGUGGGGUGAGCUGUGCUCGCGCCGCGGUUCUAUCUAGUUACGCGUUAAACCAACUAGAAUUU |
| *B2M* _L7Stop_v4 | GAGUAGCGCGAGCACAGCUAGUUUUAGAGCUAGAAAUAGCAAGUUAAAAUAAGGCUAGUCCGUUAUCAACUUGAAAAAGUGGCACCGAGUCGGUGCUCCGUGGGGUAGGCUGUGCUCGCGCCGCGGUUCUAUCUAGUUACGCGUUAAACCAACUAGAAUUU |

| Nick sgRNA | Sequence |
|---|---|
| *ATP1A1* -G3 | GAGUUCUGUAAUUCAGCAUAGUUUUAGAGCUAGAAAUAGCAAGUUAAAAUAAGGCUAGUCCGUUAUCAACUUGAAAAAGUGGCACCGAGUCGGUGCUUUU |
| *HBB* Nick 1 | CCUUGAUACCAACCUGCCCAGUUUUAGAGCUAGAAAUAGCAAGUUAAAAUAAGGCUAGUCCGUUAUCAACUUGAAAAAGUGGCACCGAGUCGGUGCUUUU |
| *HBB* Nick 2 (PE3b) | GUAACGGCAGACUUCUCUUCGUUUUAGAGCUAGAAAUAGCAAGUUAAAAUAAGGCUAGUCCGUUAUCAACUUGAAAAAGUGGCACCGAGUCGGUGCUUUU |
| B2M Nick | AGUGGAGGCGUCGCGCUGGCGUUUUAGAGCUAGAAAUAGCAAGUUAAAAUAAGGCUAGUCCGUUAUCAACUUGAAAAAGUGGCACCGAGUCGGUGCUUUU |
